# Microniches in biofilm depth are hot-spots for antibiotic resistance acquisition in response to *in situ* stress

**DOI:** 10.1101/2021.11.03.467100

**Authors:** Linda Tlili, Marie-Cécile Ploy, Sandra Da Re

## Abstract

Class 1 integrons play a major role in antibiotic resistance dissemination among Gram-negative bacteria. They are genetic platforms able to capture, exchange and express antibiotic resistance gene cassettes. The integron integrase, whose expression is regulated by the bacterial SOS response, is the key element of the integron catalyzing insertion/excision/shuffling of gene cassettes. We previously demonstrated that the basal level of integrase expression and in consequence, its activity, is increased *via* the starvation-induced stringent response in the biofilm population. However, biofilms are heterogeneous environments where bacteria are under various physiological states. Here we thus analyzed at the bacterial level, the SOS response and integrase expression within the biofilm, using confocal microscopy and flow cytometry. We showed that in the absence of exogenous stress, only a small number of bacteria (~ 1%) located in the depth of the biofilm induce the SOS-response leading to a high level of integrase expression, through both a stringent response-dependent and -independent manner. Our results thus indicate that few bacteria located in microniches of the biofilm depth undergo sufficient endogenous stress to promote the acquisition of antibiotic resistance, forming a reservoir of bacteria ready to rapidly resist antibiotic treatments.

## Introduction

In natural settings, bacteria most often live in complex communities attached to surfaces known as biofilms [1]. Biofilms are highly heterogeneous environments where bacteria experience local gradients of physical (temperature, pH) and biological (nutrients, oxygen, waste products) parameters, resulting in microniches of distinct bacterial subpopulations [2]. Early biofilm characterization studies used the biofilm as a whole, which provided global information averaging the entire biofilm bacterial population [2,3]. In the last decades, the use of fluorescent reporter genes in combination with confocal scanning laser microscopy (CLSM) has proven to be particularly useful for monitoring gene expression heterogeneity within biofilms [4–8]. Fluorescence-activated cell sorting (FACS) offered complementary information allowing to separate biofilm subpopulations to further characterize them independently [9–11]. All these techniques have helped to show that biofilms favor horizontal gene transfer [12], the main driving force for the emergence and dissemination of antibiotic resistance. One of the most striking characteristics of the biofilm is its recalcitrance towards antibiotics and host defenses, leading to treatment failure and infection recurrence[13].

Resistance integrons (RI) are low-cost structures allowing bacteria to rapidly adapt to antibiotic pressure [14,15]. They play a major role in the spread of antibiotic resistance among Gram-negative bacteria [14]. Integrons are genetic platforms able to capture, exchange and express resistance genes embedded within gene cassettes (GC) from a common promoter Pc [16]. The integrase, IntI, is the key element of the integron system, catalyzing GC insertion and excision through site-specific recombination [17]. Among the five classes of RIs identified based on the IntI amino acid sequence, class 1 RIs are the most frequently reported in antibiotic-resistant clinical isolates [18,19]. The expression of the class 1 integrase, IntI1, is controlled by the bacterial SOS response, a global regulatory network addressing DNA damages and mainly governed by the transcriptional repressor LexA [20,21]. We recently reported that in the absence of exogenous stress, the expression level of the *intI1* integron integrase, as well as that of the SOS regulon gene *sfiA*, was higher in a continuous biofilm culture than in a planktonic culture [22]. This increase depended on the RelA and SpoT proteins, the regulators of the stringent response (stress response induced upon starvation and whose effectors are the alarmones (p)ppGpp [23]). However these data were obtained from the whole biofilm and may not reflect what actually happens within its heterogeneous structure, where not all bacteria are subjected to nutrient starvation [24]. Herein, using CLSM and FACS approaches, we could refine our previous observation, and showed that actually only a very small number of bacteria located in the depth of the biofilm expressed *intI1* and *sfiA* genes at a high level in both a *relA*/*spoT*-dependent and -independent manner in the absence of exogenous stress. Most importantly our results highlight that endogenous stress in specific microniches in the depth of the biofilm would be sufficient to form a reservoir of bacteria fully ready to develop antibiotic resistance upon antibiotic treatments.

## Materials and methods

### Strains and growth conditions

Bacterial strains and plasmids used in this study are listed in Table 1. Strains were grown under planktonic or biofilm conditions at 37°C in Luria-Bertani (LB) medium supplemented with kanamycin (Km; 25μg/ml) when required.

**Table 1:**
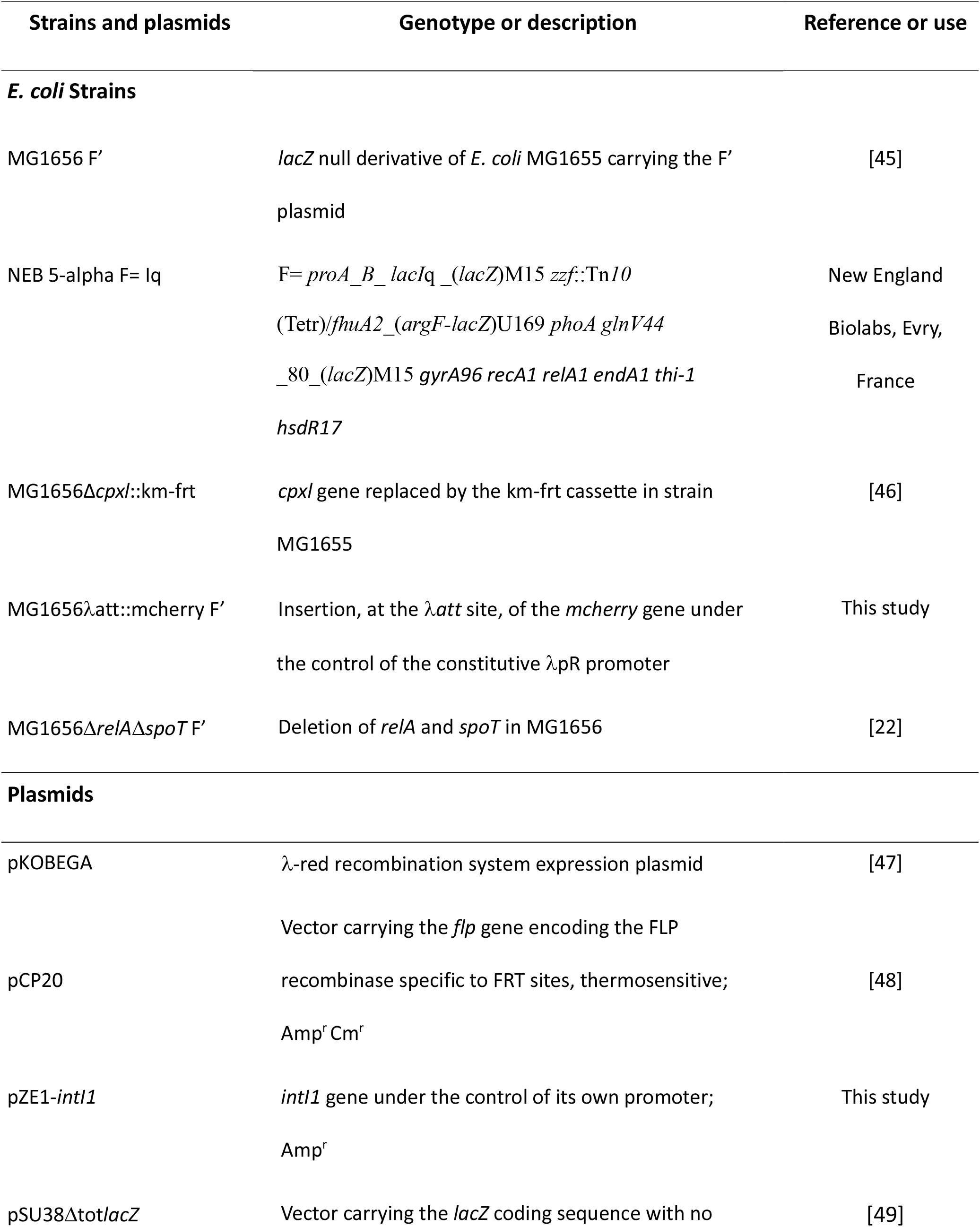
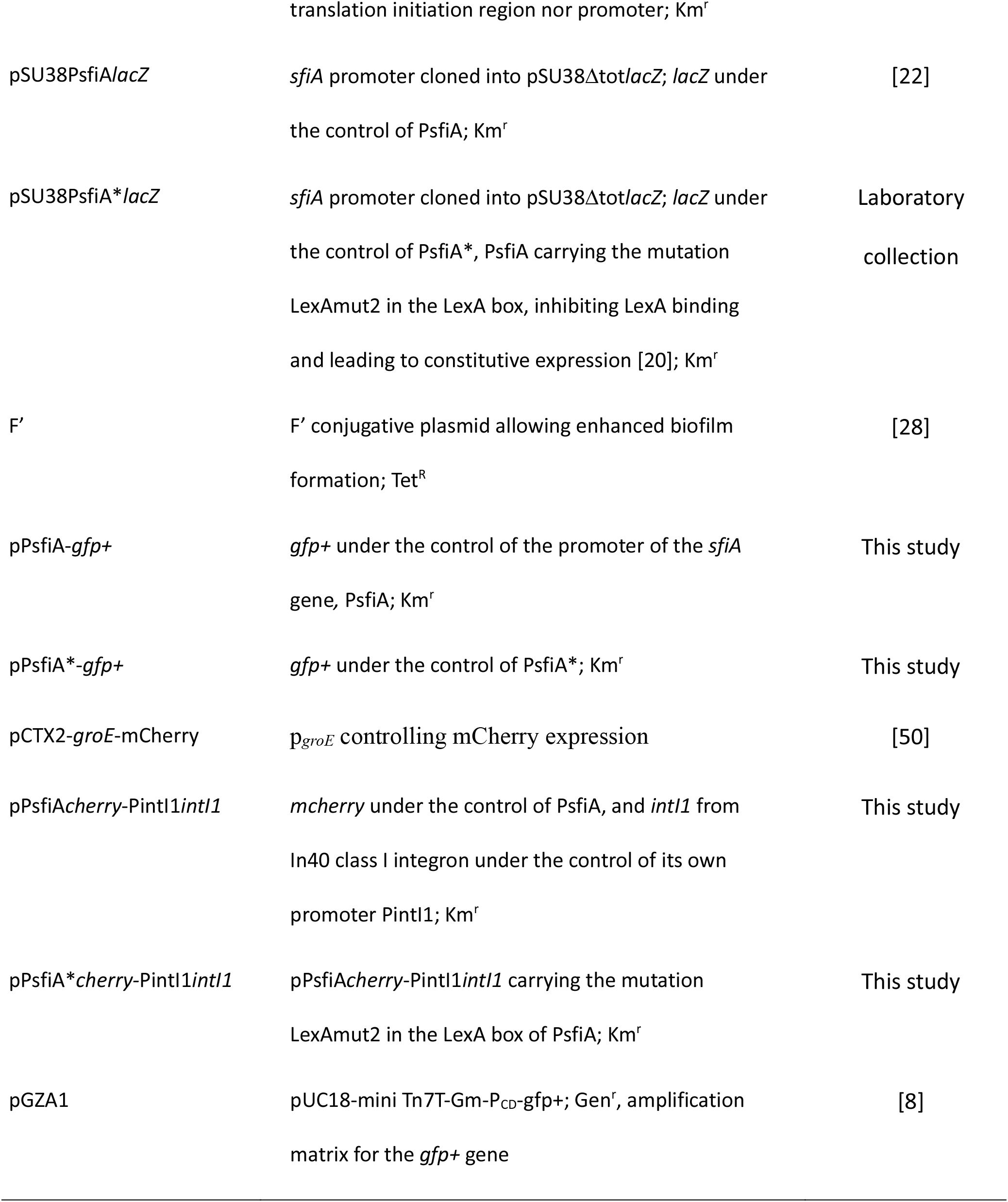
Strains and plasmids used in this study.

Biofilms in continuous culture were grown for 24 h in a glass microfermentor containing a removable spatula, as previously described [22]. Briefly, the microfermentor were inoculated by dipping the spatula for 2 min into 15 ml of over-night bacterial culture containing 1X10^9^ cells/ml, followed by a brief rinse in LB medium before insertion in the microfermentor and start of the culture. After 24h, the biofilm was resuspended by vigorously shaking the microfermentor, and the biofilm biomass was estimated by determining the optical density at 600 nm (OD600).

For planktonic culture, 100 μl of the culture used to inoculate the microfermentor was diluted in 10 ml of LB and grown for 24 h at 37°C with shaking.

### Construction of the *MG1656latt::cherry* F’ strain

The *E. coli MG1656latt::mcherry* strain, expressing red fluorescent protein (mCherry), was constructed by integrating a *km-frt-mcherry* cassette at the *E. coli λatt* site in a three-step-PCR-based approach, and removing the Km resistance gene with the FLP recombinase as previously described [25]. Primers used to construct and verify the strain, are listed in Table S1. The F’ plasmid was introduced by conjugation in this strain from the NEB5-alphaF’ strain.

### Plasmid construction

*pPsfiAgfp+* and *pPsfia* gfp+* were constructed by amplifying the *gfp+* gene (encoding a GFP variant with increased fluorescence intensity[8]) from plasmid pGZA1 (Table 1) using primers PsfiA-gfp(+)INF-5 and PsfiA-gfp(+)INF-3 and cloning the amplified fragments into plasmids pSU38PsfiA*lacZ* and pSU38PsfiA\**lacZ* respectively, digested with BamHI and HindIII, using the In-Fusion HD cloning kit (Clontech, Kusatsu, Japan) following manufacturer’s instructions. pPsfiA*cherry*-pintI1*intI1* and pPsfiA*cherry-PintI1*intI1* were constructed by amplifying first the *mcherry* gene encoding a red fluorescent protein from pCTX2-groE-mCherry using primers hindIII-mcherry and bamHI-mcherry2 and cloning the amplified fragment into pSU38PsfiA*lacZ* and pSU38PsfiA\**lacZ* respectively digested with BamHI and HindIII, yielding pPsfiA*cherry* and pPsfiA\**cherry*. Primers ter-intI-fwd-2 and inTI-pSU-rev, ter-ter-fwd-2 and ter-intI-rev-3, and pSU-ter-fwd-2 and ter-ter-rev-2, were used to amplify respectively the *intI1* gene and the lambda t0 terminator from pZE1-*intI1* (Table 1). These fragments were cloned at the XmnI site of pPsfiA*cherry* and pPsfiA\**cherry* respectively using the In-Fusion HD cloning kit (Clontech). All constructs were verified by PCR and sequencing. All primers are listed in Table S1.

### Time-lapse Microscopy with Microfluidic flow-cell

Experiments were performed with a CellASIC^®^ ONIX2 microfluidic system (www.cellasic.com). Microfluidic biofilm cultures were grown in M04S microfluidic plates (EMD Millipore, Burlington, Massachusetts, USA). After priming with culture media, the cell culture chamber was loaded with 10μl of overnight bacterial culture diluted to 5X10^7^ cells/ml with a flow pressure of 0,25 psi for 6 s. Bacteria were allowed to attach to the surface for 15min without media perfusion, before starting a laminar flow of 7μl /h (1psi) for 24h. The temperature was maintained at 37°C. Image acquisition was performed with a Carl Zeiss (Oberkochen, Germany) LSM 880 inverted confocal microscope equipped with a large incubator chamber allowing temperature control, and a 40X oil immersion objective. At least three positions were selected along the chamber for real-time imaging. Images were captured every 2h during 24h with a constant 0.4 μm Z-step and were analyzed using the IMARIS 9.6 software (oxford instruments, Tubney Wood, UK). Experiments were performed at least three times.

### Aerobic fluorescence recovery (AFR)

Resuspended 24 hours old microfermentor-biofilms were diluted in PBS 1X to a final concentration of 10^7^ bacteria/ml. Chloramphenicol (Cm) was added at a final concentration of 100 μg/ml to prevent further protein synthesis [26]. Samples were incubated at room temperature during 1h with shaking (300 rpm) to allow AFR [27].

### Determination of the proportion of red-fluorescent bacteria by flow cytometry

The resuspend biofilm after AFR was analyzed counting 10^6^ bacterial cells with a BD FACSAria III cell sorter (Becton Dickinson, Franklin Lackes, NJ, USA). Bacteria were excited with 560nm yellow green laser and red fluorescence was collected in the PE-Texas-Red channel (610/20nm). Bacteria were gated on basis of their distribution on forward *vs* side scatter plots. Red bacteria were gated based on their distribution on side scatter *vs* PE-Texas-Red fluorescence plots. The background signal was determined with a non-fluorescent bacteria biofilm. Thresholds were applied manually to determine the boundaries between the different populations and remained the same for all experiments. Data were analyzed using the FlowLogic™ software (Inivai Technologies, Mentone, Australia).

### Sorting of bacterial subpopulations

Red-fluorescent bacteria were sorted from the resuspended biofilm after AFR using a BD FACSAria III cell sorter (Becton Dickinson). Cells were collected in Nucleoprotect RNA solution (Macherey-Nagel, Dueren, Germany). Bacteria were gated as described above. The sort collection chamber was maintained at 4°C. Up to 2.10^6^ and 5.10^5^ bacteria were collected in 2 hours for respectively both intermediate- and non-fluorescent-, and super-fluorescent-biofilm populations. For the MGΔ*relA*Δ*spoT* super-fluorescent population approximatively 3.10^5^ bacteria were collected in 3 hours.

### Quantification of transcripts

The pellet of sorted bacteria were incubated in 100μl of TE (30 mM Tris-HCl, 1 mM EDTA, pH 8.0) in the presence of 100μg of lysozyme and 200μg of Proteinase K for 10 min at 37°C with shaking. Total RNA was then extracted with the RNeasy Mini Kit (Qiagen, Hilden, Germany) following manufacter’s instructions. RNA concentration, quality and integrity were evaluated with the Agilent RNA 6000 Pico kit on a Bioanalyzer. cDNA were synthesized from 100 pg of RNA and quantified using the TaqMan^®^ Fast Virus 1-Step Master Mix (Thermofisher, Waltham, Massachusetts, USA) following manufacter’s instructions with appropriate oligonucleotides and Taqman probes (Table S1). Assays were performed with a CFX96 Touch detection system (Bio-Rad^®^, Hercules, Californie, USA) using the following PCR cycling conditions: 5 min at 50°C (retro-transcription reaction (RT)), 20s at 95°C (RT enzyme inactivation) and 40 cycles at 95°C for 15s and 60°C for 45s. Assays were performed in triplicate from six independent sorting.

### Quantification of plasmid copy number

Sorted subpopulations were resuspended in 300 μl of PBS 1X and incubated at 95°C for 10 min. Debris were removed by centrifugation, 5 min at 11000 g. Concentration of the extracted DNA was measured with Qubit 4 Fluorometer (Thermofisher) and normalized to 5 pg/μl in water. Quantification of *mcherry* and *gyrB* (single gene copy on the plasmid and on the chromosomal DNA respectively) copy number was made from 25 pg of DNA using the LightCycler^®^ FastStart DNA Master HybProbe kit (Roche, Bâle, Switzerland) following manufacturer’s instructions with appropriate oligonucleotides and Taqman probes (Table S1). Assays were performed with a CFX96 Touch detection system using the following PCR cycling conditions: 10 min at 95°C followed by 40 cycles at 95°C for 30s and 60°C for 1min. Assays were performed in triplicate on each population from five independent sorting. The plasmid copy number was estimated as the ratio of *mcherry* copy number/ *gyrB* copy number.

### Statistical analysis

Statistical analysis was performed using GraphPad Prism 8 (version 8.4.3) for Windows, GraphPad Software, La Jolla California USA, www.graphpad.com. For two groups comparison a two-tailed Wilcoxon test (paired) or Mann–Whitney U-test (unpaired) test was used. Statistically significant differences were defined by a p value lower than 0.05.

## Results

### Heterogeneity of the SOS response level within the *E. coli* biofilm

To assess the spatial distribution of the bacteria inducing the SOS response and thus expressing *intI1* within the biofilm in the absence of any exogenous stress, we used a continuous microfluidic culture system coupled with confocal microscopy allowing the development of a three-dimensional biofilm in a flow chamber and its visualization in real-time. We used the *E. coli* MG1656 F’ tagged on the chromosome with the *mcherry* gene encoding a red-fluorescent protein expressed from a constitutive promoter (hereafter named MGcherry F’; Table 1). We monitored the expression of *intI1* and *sfiA* genes, both regulated by the SOS response, using transcriptional fusions between their respective promoters, PintI1 and PsfiA, and the fluorescent reporter gene, *gfp+*. The plasmid pPsfiA*-*gfp+*, in which the LexA binding region is mutated leading to constitutive expression of *gfp+*, was used as a positive control of fluorescence and SOS response. The pSUΔtot*lacZ* plasmid was used as negative control (Table 1). We knew from our previous study that in biofilm, PsfiA activity fully depends on an active SOS response, as PsfiA is inactive in a MGΔ*recA* mutant (no activator of the SOS response, RecA) and fully derepressed in a MGΔ*lexA* mutant (no repressor of the SOS response, LexA) [22].

Biofilms of MGcherry F’ carrying one of the plasmids were grown for 24h in LB at 37°C. As expected, the CLSM images showed that the MGcherry F’/pSUΔtot biofilm contained only red bacteria and the MGcherry F’/pPsfiA*-*gfp+* biofilm contained bacteria that were both red and bright green (Fig. 1a and 1b). As seen on Fig. 1c and 1d, the MGcherry F’/pPsfiA-*gfp+* biofilm contained mainly red bacteria and very few both red and green bacteria. The 1D dot plot representing the distribution of bacteria as a function of their spatial localization further showed an uneven distribution of green bacteria on the z-axis of the 3D image of the biofilm compared to the x- and y-axes (Fig. 1g, 1e and 1f respectively). These results indicated that in the absence of exogenous stress, the *PsfiA* promoter is activated (green bacteria) in a minority of bacteria preferentially located in the depth of the biofilm.

**Fig. 1:**
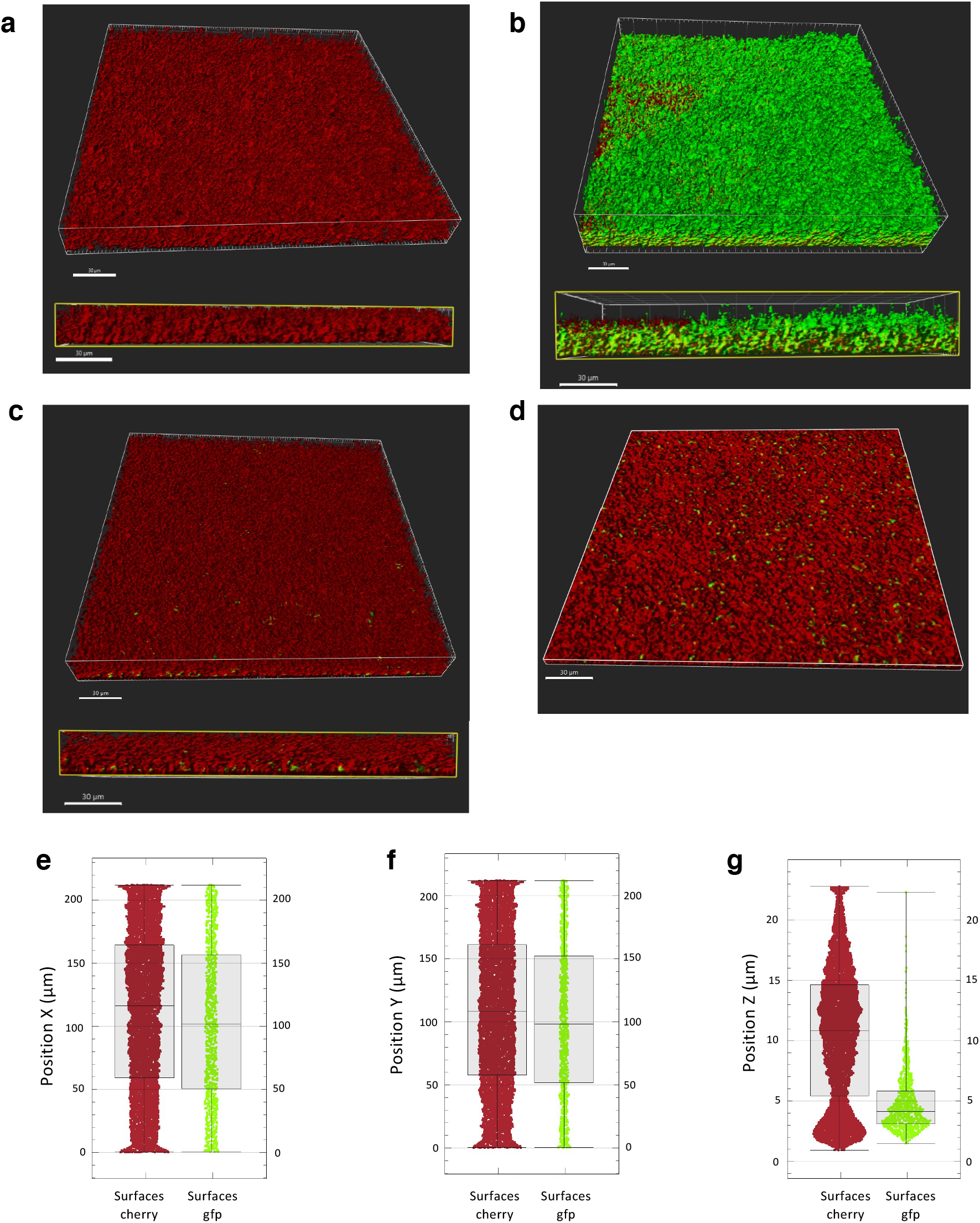
*sfiA* expression is heterogeneous and localized within the biofilm. **a**, **b**, **c**, CLSM images and section of the XZ planes of respectively *MGcherry* F’/pSUDtot, *MGcherry* F’/PsfiA*-*gfp+* and *MGcherry* F’/PsfiA-*gfp+* biofilms grown for 24h at 37°C in microfluidic flow-cell chamber. **d**, XY plan section of the *MGcherry* F’/pPsfiA-*gfp+* biofilm image (c) at 5 μm depth. z-stacks were generated using a Zeiss LSM 880 microscope and processed with Imaris 9.6 software (Bitplane). Scale bar represents 30 μm. **e**, **f**, **g**, 1D plot representing the total number of green and red fluorescent bacteria in the biofilm as a function of their X-position, Y-position and Z-position respectively. Boxplots are designed as median, the lower quartile (Q1) and the upper quartile (Q3). Whiskers are extended to the extreme data points. All images are representative of one position among 10 taken for each condition. Experiments were done in triplicate.

We then estimated the proportion of these bacteria in the biofilm, using a FACS approach, which allowed to analyze each bacterium separately. To obtain enough biomass for FACS analysis, we grew our biofilms in a continuous culture in microfermentor [28]. Surprisingly, we detected no red fluorescence from the MGcherry F’/pPsfiA*-*gfp+* biofilm, and only 87% of the bacteria in the biofilm emitted green fluorescence, which was of very low intensity. We postulated that this lack or low level of fluorescence was related to the biofilm model itself. Indeed, as the microfermentor-biofilm is thicker and larger than the microfluidic flow chamber one, the oxygen concentration might be more limiting, therefore impairing the maturation of the GFP+ and mCherry fluorescent proteins [27].

To complete the protein maturation to a fully fluorescent form, we added a step of AFR after biofilm growth [27]. The resuspended microfermentor-biofilm was incubated for one hour at room temperature with shaking in the presence of chloramphenicol to prevent further protein synthesis [26]. This method allowed full recovery of red fluorescence (mCherry) with high fluorescence emission intensity. However, for the green fluorescence, the fluorescence intensity remained very low and too close to the background signal to be properly quantified. Therefore, the plasmid model was adjusted accordingly. We constructed PsfiA and PsfiA* transcriptional fusions with *mcherry* as the reporter gene (to follow PsfiA activity in microfermentor-biofilms by FACS). And as we were unable to visualize by CLSM *intI1* expression in the biofilm of MGcherry F’/pintI1-*gfp+*, probably due to the low level of *intI1* expression, even when the *intI1* gene is derepressed [22], we also added on the new plasmid, the *intI1* gene under the control of its own promoter PintI1, to subsequently estimate integron integrase expression by RT-qPCR (see below). These plasmids were transformed in the strain MG1656 F’, leading respectively to strains MG F’/pPsfiA*cherry*-PintI1*intI1* and MG F’/pPsfiA\**cherry*-PintI1*intI1* (Table 1).

After one hour of AFR, 99 % of the bacteria of a 24h-old microfermentor-biofilm of the positive control MG F’/pPsfia\**cherry-*PintI1*intI1* were bright red fluorescent (Fig. 2b). For MG F’/pPsfiA*cherry-*PintI1*intI1*, FACS dot-plot analysis showed that, as observed with the microfluidic chamber, only few bacteria expressed fluorescence in the biofilm. This approach further showed that this minority of red bacteria (3.3 ± 1.8 %) is clearly divided into two populations: one (2.3 ± 1.2 %) expressing red fluorescence at an intermediate level, the other (1 ± 0.7 %) at a high level (Fig. 2c). The latter showed red fluorescence intensity equivalent to that expressed from the constitutive promoter PsfiA*, indicating that in this population the SOS response is strongly induced, leading to full activity of the PsfiA promoter. These two red populations will be named hereafter “intermediate-” and “super-fluorescent-populations” respectively.

**Fig. 2.**
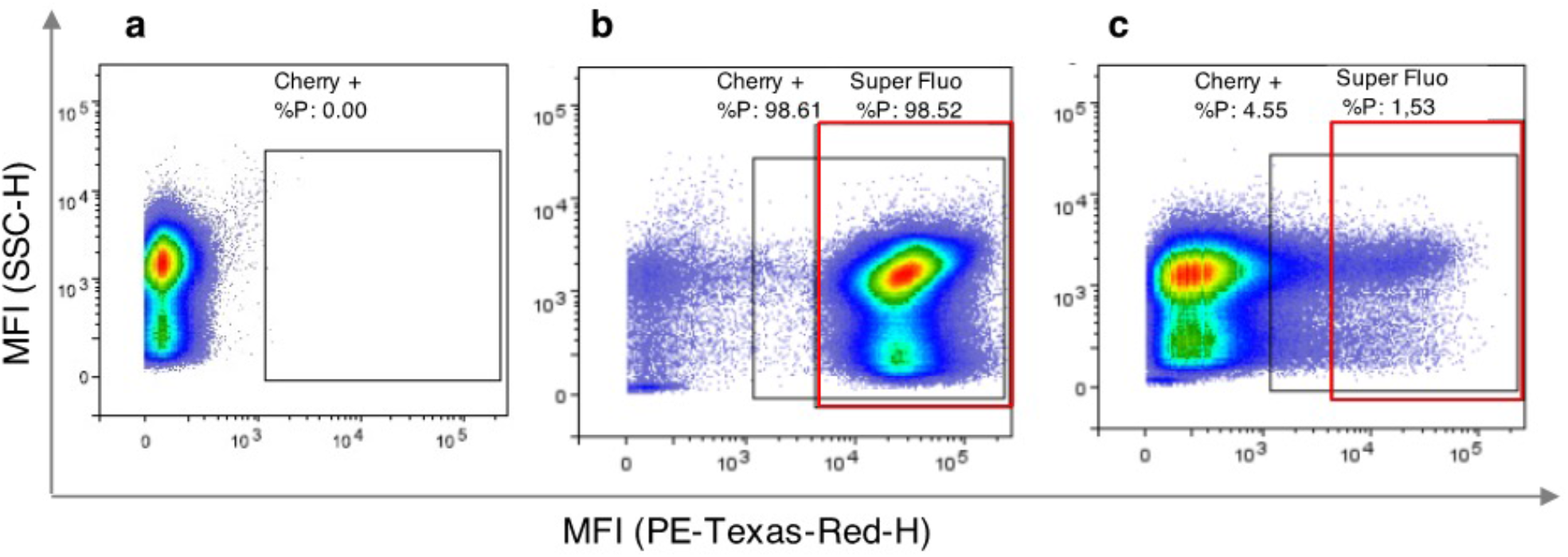
Quantification of cells expressing *sfiA* in biofilm. Flow cytometry dot-plot analyses of a 24h-old biofilm of strain (**a**) MG F’/pSUΔtot, (**b**) MG F’/pPsfiA\**cherry*-PintI1*intI1* and (**c**) MG F’/pPsfiA*cherry*-PintI1*intI1* grown in continuous culture in microfermentor. Counting was performed after one hour of aerobic fluorescence recovery at room temperature. The black square delimits the cherry-positive population (cherry +) based on the non - fluorescent strain MG F’/pSUΔtot (panel a), and the red rectangle defines the cherry-super-fluorescent population (super Fluo) based on derepressed cherry expression in strain MG F’/ pPsfiA\**cherry*-PintI1*intI1* (panel b). The data represent one representative experimental replicate.

### Differential Integron integrase expression within biofilm subpopulations

As we have previously shown that basal-expression of the *intI1* and *sfiA* genes was higher in biofilm than in planktonic culture without any exogenous stress [22], we hypothesized that the expression of *intI1* and *sfiA* would happen within the same subpopulations in the biofilm. We therefore sorted by FACS the three non-fluorescent, intermediate and super-fluorescent subpopulations of the MG F’/pPsfiA*cherry-*PintI1*intI1* biofilm, in order to quantify in each of them the transcript levels of *intI1* and *mcherry* genes (expressed from promoters PintI1 and PsfiA respectively) by RT-qPCR. First, we confirmed that no degradation of the bacterial RNA or changes in the level of *mcherry* expression occurred in the initial culture (*i.e*. re-suspended biofilm during the various steps of the experiment (AFR + sorting)) (Fig. S1).

Transcript levels were higher in the intermediate population (1.9- and 1.7-fold for *cherry* and *intI1* respectively) and in the super-fluorescent population (64- and 22-fold for *cherry* and *intI1* respectively) than in the non-fluorescent population (Fig. 3). As the *sfiA* gene is present in the chromosome of our MG F’/pPsfiA*cherry*-PintI1*intI1* strain, we therefore quantified *sfiA* transcripts to verify that its expression was also induced in the fluorescent populations of the biofilm. Surprisingly, the chromosomal *sfiA* transcript level was significantly increased but only in the super-fluorescent population and at a much lower level than that of the plasmid genes *intI1* or *mcherry* (Fig. 3). Altogether, these results suggested that the strong increase in *mcherry* and *intI1* transcripts may be influenced by variations in plasmid copy number in the biofilm subpopulations, in addition to the induction of the SOS response.

**Fig. 3.**
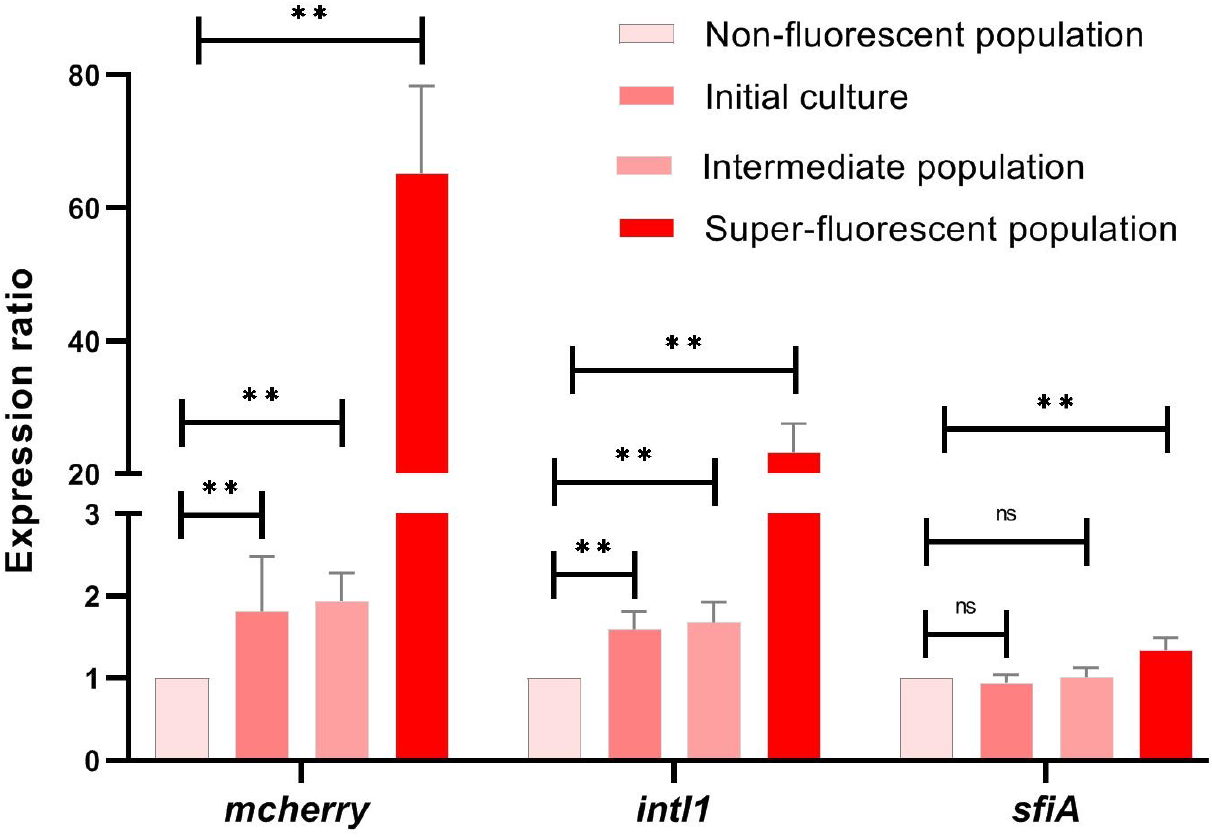
Differential expression of *mcherry, intI1* and *sfiA* in biofilm subpopulations. Expression ratio of *mcherry, intI1* and *sfiA* transcripts from initial culture (dark pink) and sorted sub-populations (non-fluorescent (light pink), intermediate (medium pink) and super-fluorescent (red)) of a 24-old biofilm of MG F’/ pPsfiA*cherry*-PintI1*intI1* grown in microfermentor. Relative quantification of *mcherry, intI1* and *sfiA* was performed using the 2^-ΔΔCt^ method, the transcript number of each gene was normalized to the endogenous *gyrB* gene and calibrated to the non-fluorescent population. Data are the average of transcripts levels measured from 6 independent sorting experiments. Errors bars indicate the SD. Asterisks indicate significant difference: ** p<0.01; ns: non-significant (Wilcoxon test).

### Influence of biofilm microniches on plasmid copy number

To answer the above questioning, we quantified the plasmid copy number in the three populations of the MG F’/pPsfiA*cherry-*PintI1*intI1* biofilm. As shown on Fig. 4a, the plasmid copy number increased by 1.9- and 14.2-fold respectively in intermediate and super-fluorescent populations compared to the non-fluorescent population. We thus also quantified the transcript level of the kanamycin resistance gene, *aph(3’)-IIa*, carried on the same plasmid as the *mcherry* and *intI1* genes, and constitutively expressed. The *aph(3’)-IIa* transcript level should only reflect variations in the plasmid copy number. We observed that the *aph(3’)-IIa* expression level increased in both the intermediate- and super-fluorescent-populations, in proportions similar to plasmid copy number (Fig. 4b). Therefore, from now on, we normalized the transcript levels of *mcherry* and *intI1* genes by that of *aph(3’)-IIa* to estimate the effect of the SOS response induction on PsfiA and PintI1 activity. As shown on Fig. 4c, the activation of PsfiA and PintI1 occurred only in the super-fluorescent population (by 4.1- and 1.6-fold factor respectively), in agreement with the above observations for chromosomal *sfiA* transcript levels (Fig. 3). Overall, our results show that, in the biofilm, there are two bacterial populations that express SOS-inducible genes at different levels: one expressing these genes at their low basal non-induced level (non-fluorescent and intermediate-populations), and one at a high de-repressed level (super-fluorescent population).

**Fig. 4:**
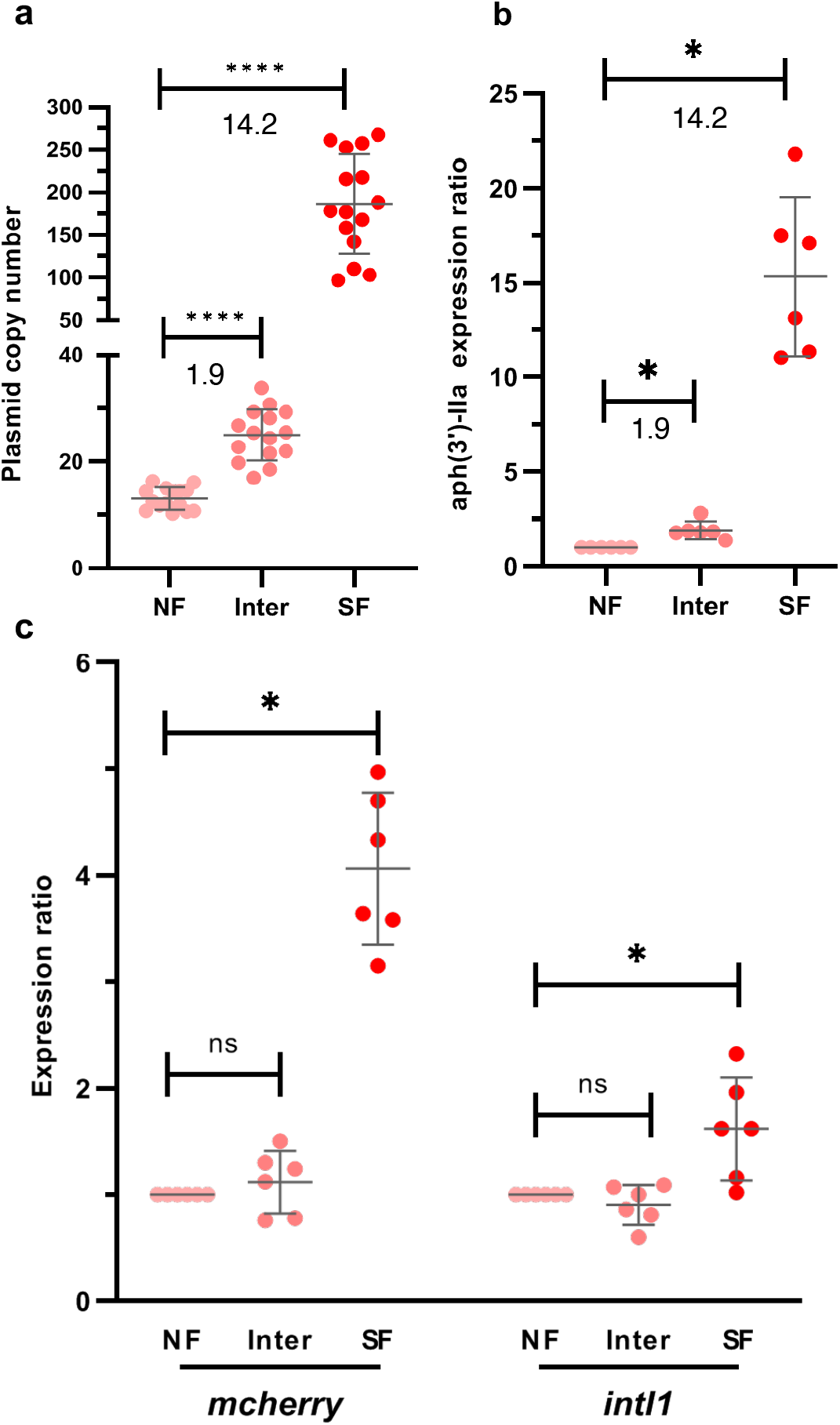
Impact of the variation in plasmid copy number in the biofilm subpopulations on *mcherry* and *intI1* expression level. **a**, The plasmid copy number was estimated by calculating the ratio of gene copies number of plasmidic *mcherry* over chromosomal *gyrB* for each sorted sub-populations from a 24h-old biofilm of MG F’/ pPsfiA*cherry*-PintI1*intI1* strain grown in microfermentor. **b**, Expression ratio of *aph(3)-IIa* transcripts from sorted sub-populations NF, inter and super + (same biofilm as in a). Relative quantification of *aph(3’)-II* (gene encoding kanamycin resistance carried on the same plasmid as *mcherry)* was performed using the 2^-ΔΔCt^ method, with normalization to the endogenous *gyrB* gene and calibration to the non-fluorescent population. **c**, Expression ratio of *mcherry* and *intI1* transcripts in sorted sub-populations, NF, inter and super +. Relative quantification of *mcherry* and *intI1* was performed using the 2^-ΔΔCt^ method, with normalization to the *aph(3)-IIa* gene and calibration to the non-fluorescent population. NF (non-fluorescent; light pink), inter (intermediate; medium pink) and SF (super-fluorescent; red) sub-populations. Data are the average of gene copies number or transcripts levels measured from 6 independent sorting experiments: Average and standard deviation are shown as black lines (a and c), errors bars indicate the SD (b). Asterisks indicate significant difference: * p<0.05; **** p<0.0001; ns: non-significant (Wilcoxon test).

### Induction of SOS response in the super-fluorescent population is only partially dependent on the stringent response

We previously reported that the basal expression *of sfiA* was 2.2-fold higher in biofilm than in planktonic culture and that this increase was dependent on the stringent response, which is induced by the RelA and SpoT proteins [22]. To determine whether the super-fluorescent population was specific of the biofilm and dependent on the stringent response, we analyzed by FACS a 24h-old planktonic culture of MG F’/pPsfiA*cherry*-PintI1*intI1*, as well as a 24h-old microfermentor-biofilm of a mutant unable to induce the stringent response, MGΔ*relA*Δ*spoT* F’/pPsfiA*cherry*-PintI1*intI1* (Table 1). Both cultures contained a super-fluorescent population but in significantly lower proportion than in the MG F’/pPsfiA*cherry*-PintI1*intI1* biofilm (Fig. 5a and 5b). These results suggested that more bacteria activated the SOS-inducible promoters PintI1 and PsfiA in the biofilm than in the planktonic culture, and that in the biofilm about half of the super-fluorescent population could induce the SOS response independently of the stringent response. The next question was are the expression levels of *intI1, mcherry* and *sfiA* similar in all three super-fluorescent populations *i.e*. from planktonic and both biofilms (MG F’ and Δ*relA*Δ*spoT* mutant) cultures?

**Fig. 5:**
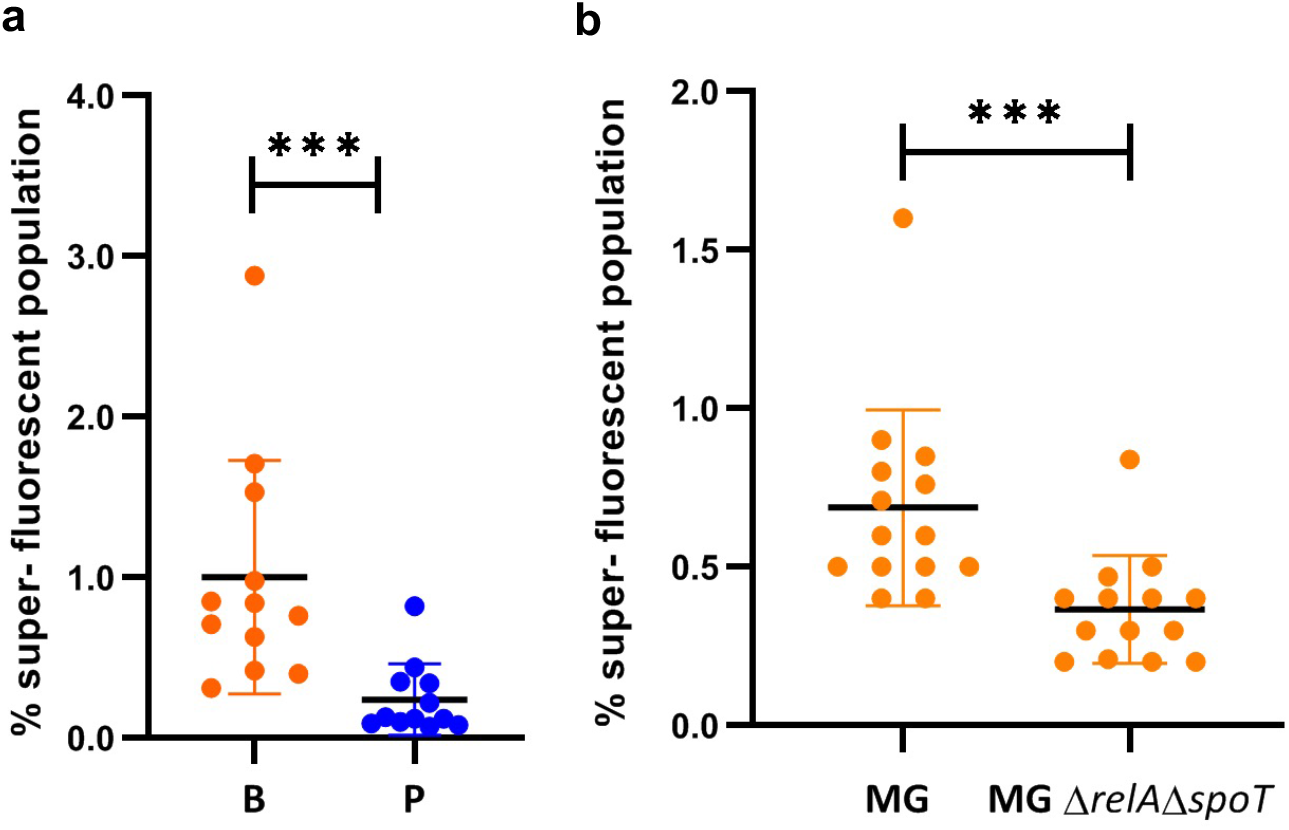
Comparison of cherry expression level in biofilm and planktonic conditions, and effect of the stringent response on mcherry expression in biofilm. Proportion (%) of cherry-super-fluorescent populations in (**a**) biofilm (B, orange) and planktonic (P, blue) cultures of the MG *F’/pPsfiAcherry-PintI1intI1 strain;* (**b**) in biofilm of MG F’ and MGΔ*relA*Δ*spoT* F’ strains carrying plasmid pPsfiA*cherry*-PintI1*intI1*. Data are the average of at least 12 independent cultures. Asterisks indicate significant difference between biofilm and planktonic conditions (a) or between MG and MGΔ*relA*Δ*spoT* (b); *** p<0.001 (Mann-Whitney U-test (a) or Wilcoxon test (b)). Average and standard deviation are shown as black lines.

### Complexity of the regulation of *intI1* expression by the stringent response in biofilm

The amount of super-fluorescent bacteria was too low in the planktonic culture to be sorted in a timescale suitable with our experiment. However, we were able to sort non-fluorescent and super-fluorescent populations from a 24h-old MGΔ*relA*Δ*spoT* F’/pPsfiA*cherry*-PintI1*intI1* biofilms by FACS in sufficient amounts to quantify *intI1*, *mcherry* and chromosomal *sfiA* transcripts by RT-qPCR. Transcript levels from plasmidic *mcherry* and *intI1* genes were normalized to *aph(3’)-IIa*, the plasmid copy number being also higher in the super-fluorescent MGΔ*relA*Δ*spoT* F’/pPsfiA*cherry*-PintI1*intI1* population (Fig. S2).

In the Δ*relA*Δ*spoT* mutant background, the transcript level of chromosomal *sfiA* was not significantly higher in the super-fluorescent population compared to the non-fluorescent population, contrary to what was observed in the wild type (WT) MG F’ background (Fig. 6a), suggesting that the significant increase of *sfiA* expression in MG F’/pPsfiA*cherry*-PintI1*intI1* super-fluorescent population depended on the stringent response. However, the transcript levels of the plasmid genes, *mcherry* and *intI1*, were significantly higher in the MGΔ*relA*Δ*spoT* super-fluorescent population than in the non-fluorescent population (Fig. 6b). Nevertheless, *intI1* expression ratio was still higher in the super-fluorescent population of the WT MG F’ background than in the mutant one (Fig. 6b). The trend was the same for the *mcherry* gene, although the difference of expression between the WT and mutant background super-fluorescent populations did not appear to be significant (Fig. 6b). Altogether these results indicated that for the *mcherry* and *intI1* genes, the stringent response participates but is not the only stress to be involved in the SOS response induction in the super-fluorescent population of the biofilm.

**Fig. 6:**
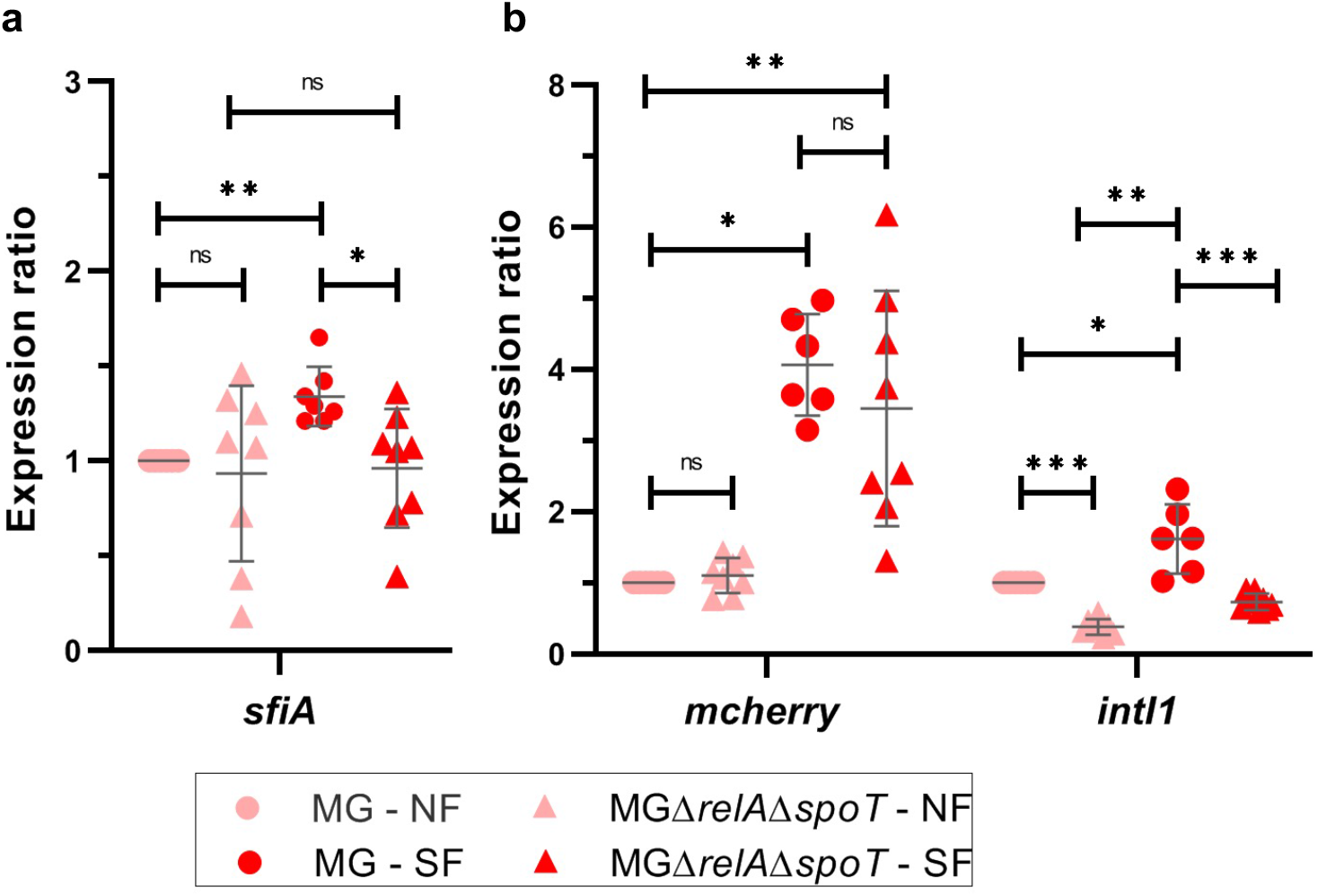
The stringent response induces the expression of sfiA, mcherry and intI1 in the biofilm super-fluorescent subpopulations. Expression ratio of (a) *sfiA*, (**b**) *mcherry* and *intI1* transcripts number in non-fluorescent (NF; light pink) and super-fluorescent (SF; red) sorted sub-populations of 24-old biofilms of MG F’ (circle) and MGΔ*relA*Δ*spoT* F’/ (triangle) strains carrying pPsfiA*cherry*-PintI1*intI1*. Relative quantification of *mcherry, intI1* and *sfiA* was performed using the 2^-ΔΔCt^ method. The number of *mcherry* and *intI1* transcripts was normalized to that of *Kana* gene (constitutively expressed from the plasmid) to take in to account plasmid number variation; the transcripts number of chromosomal *sfiA* gene was normalized to the endogenous *gyrB* gene. The calibrator was MG F’/ pPsfiA*cherry*-PintI1*intI1* non-fluorescent sub-population. Data are the average of transcripts levels measured from at least 6 independent sorting experiments. *** p<0.001; ** p<0.01; ns: non-significant; Mann-Whitney U-test (comparison of WT MG F’ vs MGΔ*relA*Δ*spoT* F’) or Wilcoxon test (comparison of NF and SF populations of the same biofilm). Average and standard deviation are shown as black lines.

Surprisingly, although the level of *mcherry* expression was the same in the non-fluorescent WT and mutant populations, as observed for chromosomal *sfiA* expression (Fig. 6), *intI1* expression was significantly lower in the non-fluorescent mutant population than in the WT population (Fig. 6b). These results suggested that in the non-fluorescent population, the stringent response activates *intI1* expression in a SOS-independent manner.

## Discussion

Bacterial biofilms are highly heterogeneous environments where bacteria experience various stress [2]. Working with the total biofilm population forces one to average and minimize the observed effects, or even miss some of them, when they occur in a small number of bacteria, as shown here. Indeed, we previously showed that the expression level of *intI1* and *sfiA* genes in the absence of exogenous stress (basal expression) was higher, but not fully de-repressed, in biofilm than in planktonic culture [22]. Here, analyzing the biofilm at the bacterial level, we showed that actually only few bacteria (around 1%) located within the depth of the biofilm activated the two promoters, PintI1 and PsfiA, at a strong level. FACS analyzes were more sensitive than CLSM identifying 3 populations with different level of fluorescence (non-, intermediate- and super-fluorescent, representing in average respectively 97%, 2% and 1% of the bacteria in the biofilm; Fig. 2). However, contrary to the super-fluorescent population, in the intermediate population, the fluorescence did not correlate with increased PsfiA and PintI1 activities, it only reflected an increase in the copy number of the ColE1-type plasmid carrying the PsfiA-*mcherry* transcriptional reporter and the integron integrase PintI1-*intI1* (Fig. 4). It has been shown that starvation/limitation of specific amino acids (Thr, Arg and Leu among those tested) led to a rise in the copy number of ColE1-type plasmids in both *relA+* and *relA-E. coli* strains. This rise was even greater in the *relA-* strain than in the *relA+* strain (as observed in this study, Fig. 4a and S2) for starvation of specific amino acids (Ile, Thr and His) [29]. These observations therefore confirmed that in the biofilm, bacteria undergo amino acid limitations, and further indicated that the stronger the limitation, the higher the plasmid copy number. Indeed, the super-fluorescent population which has the highest plasmid copy number is the one that undergoes a sufficiently strong amino acids limitation to induce the stringent response, which in turn would activate the SOS response [22,30]. FACS and RT-qPCR analyses also suggested the existence of other regulation pathways different from the stringent response, that could induce the SOS response, activating both PsfiA and PintI1 in the super-fluorescent population of the MGΔ*relA*Δ*spoT* mutant biofilm (Fig. 5 and 6b). Biofilms of *E. coli* Δ*relA*Δ*spoT* mutants have been found to have reduced catalase activity (*i.e*. lower oxidative stress defense) and elevated reactive oxygen species (ROS) OH• levels [31]. ROS are known to induce the SOS response [32]. However the super-fluorescent bacteria are mainly located in the depth of the biofilm (Fig. 1) where the oxygen concentration is commonly low [2]. Therefore, the likelihood that oxidative stress *via* ROS production is one of the signals that induce the SOS response in the super-fluorescent population of Δ*relA*Δ*spoT* mutant biofilm does not seem very likely, but would need to be investigated. Unfortunately, we could not verify this hypothesis as were not able to construct an MGcherryΔ*relA*Δ*spoT* F’ mutant strain to monitor and localize the PsfiA-*gfp+* expression in flow-cell, as we did with the WT strain (Fig. 1). Another stress inducing the SOS response in the biofilm, could be phosphate starvation, which has also been found to induce the LexA regulon [33].

We were surprised to find that the level of *intI1* expression in the non-fluorescent Δ*relA*Δ*spoT* population was significantly lower than in the non-fluorescent WT MG F’ population (Fig. 6b). It indeed suggested that the basal activity of PintI1 is reduced in the non-fluorescent population in the absence of (p)ppGpp (alarmones produced by RelA and SpoT upon nutrient starvation). Since this effect was not observed for the PsfiA promoter, it appears to be specific to PintI1 and independent of the SOS response. Interestingly, Strugeon *et al*. have shown that in biofilm, the expression of PintI1* (PintI1 promoter with a mutated LexA-binding-site) is regulated by the Lon protease, which was not the case with WT PintI1 [22]. The Lon protease is known to be activated by the polyphosphate (poly-P) that accumulates upon the stringent response induction. In the model proposed by Strugeon *et al*., the poly-P-Lon complex would control the stability of an unidentified regulator of PintI1 (repressor) that would only act when the PintI1 promoter is released from LexA. Thus, one could imagine that in the non-fluorescent population of the Δ*relA*Δ*spoT* mutant biofilm, the basal level of poly-P-Lon is no longer sufficient to ensure the degradation of the inhibitor of the derepressed PintI1 promoter. In this case, the basal activity of PintI1 would decrease due to the absence of (p)ppGpp and the lower level of poly-P-Lon. Another hypothesis to explain this decrease in basal PintI1 activity in the Δ*relA*Δ*spoT* non-fluorescent population would be a direct effect on the transcription. Indeed, (p)ppGpp regulates positively or negatively transcription initiation from specific promoters by binding directly to RNA polymerase (RNAP) at two sites [34,35]. In these (p)ppGpp regulated promoters, a discriminator region (region between the conserved −10 promoter element and the transcription start site (TSS)) with more A+T rich content was linked to activated promoters [35,36]. PintI1 shares some characteristics of (p)ppGpp activated promoters (conserved bases in the −10 region, T and A bases in the discriminator region and TSS [35,37]). It is tempting to assume that the *intI1* promoter basal activity in the biofilm might be stimulated by (p)ppGpp, but this will need to be confirmed.

Overall, our results showed that the regulation of integron integrase expression in biofilm is complex. There are two main populations with different characteristics in the biofilm: the vast majority of the bacteria expressing the SOS-regulon gene *sfiA* and *intI1* at a basal level, which for *intI1* expression seems to depend on (p)ppGpp; and a minor population (1%), located in the depth of the biofilm that faces amino acid limitations leading to the induction of the stringent response, as well as other limitations (phosphate ?), all being endogenous stresses inducing the activation of SOS-regulated promoters in this population (Fig. S3). As a result, this population would be able to acquire antibiotic resistance genes *via* the expression of *IntI1*.

Dorr *et al*. showed that the formation of *E. coli* persisters (subpopulation of bacteria (usually less than 1%) surviving in presence of high bactericidal antibiotic pressure without an increase in MIC [38,39]) was induced by the SOS response after fluoroquinolone treatment [40]. Bernier *et al*. demonstrated that the tolerance of *E. coli* biofilm to ofloxacin depended on starvation and SOS response [41]. Intriguingly, our super-fluorescent-biofilm population shares several characteristics describing triggered persistence: i) it is induced upon amino acids limitation (via the stringent response) and possibly other nutrients limitations, ii) it represents a minor proportion (around 1%) of the biofilm and iii) both states (super-fluorescence/SOS induction and persistence) are transient. Indeed, persistent bacteria, when treated with bactericidal antibiotics, will regrow when diluted in an antibiotic-free media, their progeny being as sensitive to drugs as the parental population. And, in our case, when grown again in microfermentor, sorted super-fluorescent bacteria of the MG F’/PsfiA*cherry*-PintI1*intI1* biofilm gave rise to a new biofilm containing the same sub-populations as in the initial biofilm (Fig. S4). It is tempting to infer that the super-fluorescent-biofilm population could be a persister population induced by endogenous stresses of the biofilm. If so, it would mean that in the biofilm, the super-fluorescent population, which expresses the SOS response at high level, would be both tolerant to antibiotics and ready to express antibiotic resistance through gene cassettes acquisition or shuffling via the integron integrase. Furthermore, the activation of the SOS response in the super-fluorescent population could also induce error-prone DNA polymerases leading to mutations involved in antibiotic resistance (Fig. S3). In the case of antibiotic treatment, these features would allow resistant bacteria to be readily selected for. It was indeed recently shown that persisters of *E. coli* after fluoroquinolone treatment displayed enhanced resistance toward several antibiotics following recovery from treatment, and this was linked to the SOS response and the error-prone polymerase V [42].

It has been long supposed in clinical practice, that the main characteristic hampering biofilm treatment was their thick-layered structure, limiting the penetration of antibiotics, thus exposing them to sub-MIC concentrations that would facilitate the selection of antimicrobial resistance. Nowadays, it is clear that biofilm endurance is multifactorial [13,43]. It was shown in a *P. aeruginosa* biofilm that some bacteria undergo endogenous oxidative stress leading to double-strand DNA breaks. The mechanisms involved in repair of these DNA breaks, which were SOS-response-independent, generated genetic variations within the population, conferring various selective advantages, in particular mutations that increased the emergence of antibiotic resistant bacteria that could be selected for when biofilms were exposed to antibiotic treatment [44]. Here we demonstrate, that in addition to its structural advantage, the biofilm lifestyle promotes levels of local endogenous stresses, even in the absence of antibiotic pressure, that give rise to a small population of bacteria expressing at high-level, genes regulated by the SOS response. This population which shows most of the characteristics required for tolerance to antibiotics and for antibiotic resistance acquisition, could constitute a reservoir of bacteria fully ready to rapidly resist antibiotic treatment, being the cause of recurrent infections and impacting the management of nosocomial infections and antibiotic treatments.

## Acknowledgements

We thank the engineers of the technology facility BISCEm Inserm US042, CNRS UMS 2015, Claire Carrion and Catherine Ouk for their precious help with the confocal microscopy and flow cytometry experiments respectively. We are grateful to Yohann Lacotte and Cécile Pasternak for helpful discussions and critical reading of the manuscript, and to Ana Catalina Hernandez for her fruitful help and discussions for the cytometry experiments and analyses, and critical reading of the manuscript. This work was supported by grants from Ministère de l’Enseignement Supérieur et de la Recherche, Institut National de la Santé et de la Recherche Médicale (Inserm) and Fondation pour la Recherche Médicale (FRM DEQ20150331742 to MC Ploy). The funders had no role in study design, data collection and analysis, and the decision to submit this work for publication.

