## Supplemental files for "Microniches in biofilm depth are hot-spots for antibiotic resistance acquisition in response to *in situ* stress"

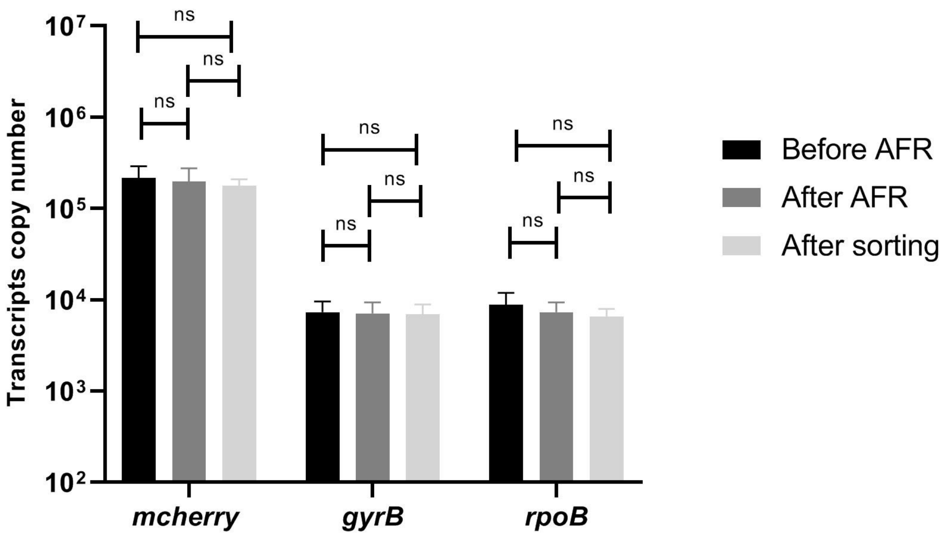


**Fig. S1: Stability of RNA**

Transcript numbers of *mcherry*, *gyrB* and *rpoB* genes were quantified by RT-qPCR in the initial population (re-suspended 24h biofilm) of MG F'/ pPsfiA*cherry*-PintI1*intI1* strain grown in microfermentor, before AFR (black), after AFR (dark grey) and after sorting (light grey). Data are the average of transcripts levels measured from 6 independent sorting experiments. Errors bars indicate the SD. ns: non-significant (Wilcoxon test). AFR: aerobic fluorescence recovery.

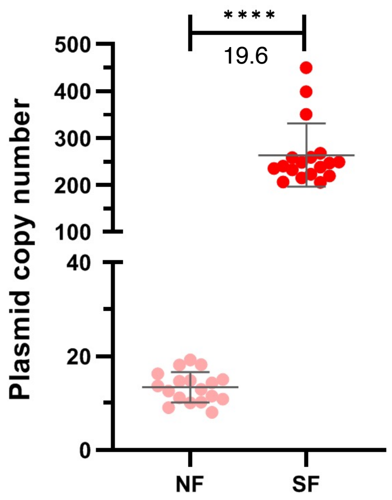


**Fig. S2: Plasmid copy number in mutant MGΔ*relA*Δ*spoT***

The plasmid copy number was estimated by calculating the ratio of gene copies number of plasmidic *mcherry* over chromosomal *gyrB* for each sorted sub-populations from a 24h-old biofilm of MGΔ*relA*Δ*spoT* F'/ pPsfiA*cherry*-PintI1*intI1* strain grown in microfermentor.

NF (non fluorescent; light pink), and SF (super-fluorescent; red) sub-populations. Data are the average of 6 independent sorting experiments. Asterisks indicate significant difference: **** $p<0.0001$ (Wilcoxon test). Average and standard deviation are shown as black lines.

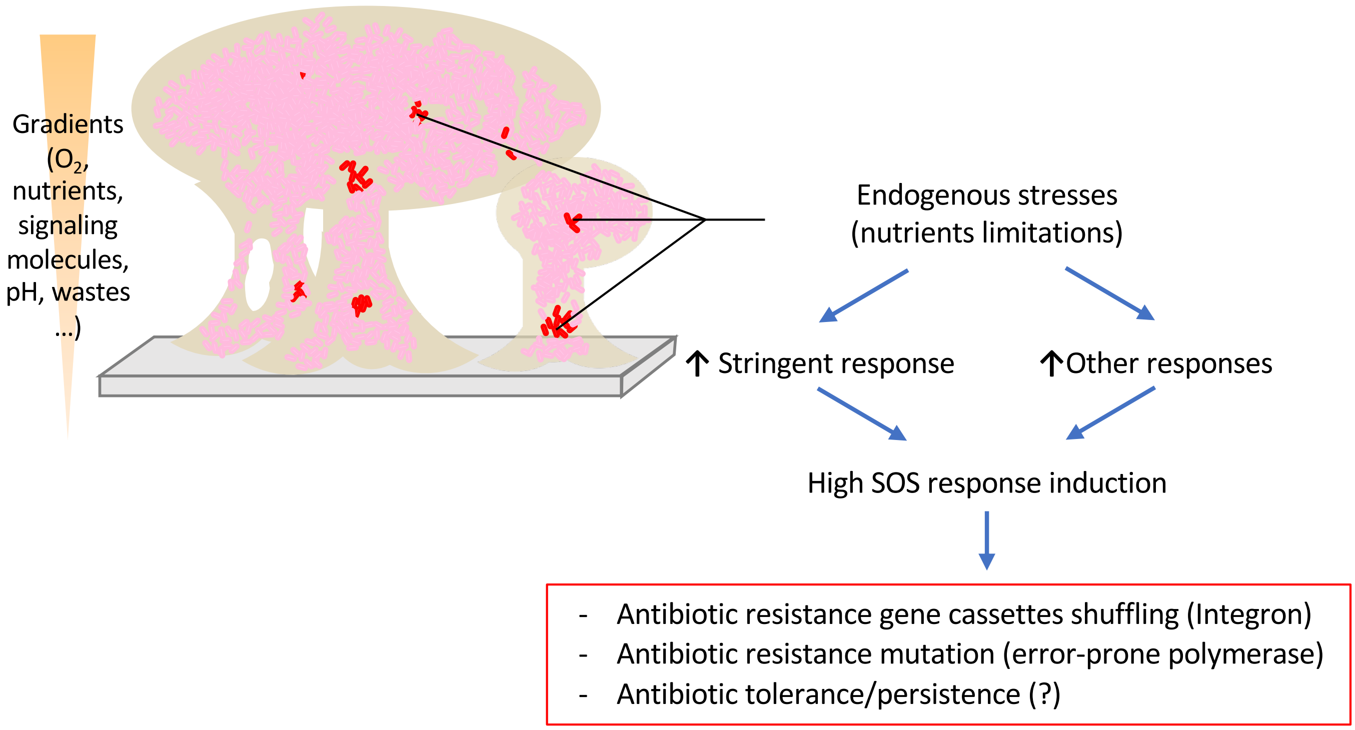


**Fig. S3: Heterogeneous expression of *sfiA* and *intI1* in the biofilm**

In the biofilm, the vast majority of bacteria (non-fluorescent and intermediate populations, 99% of the bacteria, pink bacteria) express the *sfiA* and *intI1* genes at a low basal level. Only few bacteria in the biofilm (super-fluorescent population, 1%, red bacteria) experience a level of endogenous stress (various nutrients limitations) high enough to induce the SOS response in a stringent response-dependent and stringent response-independent manner. This induction thus allows for high-level expression of genes regulated by the SOS response, leading to various outcomes favoring the acquisition/expression of antibiotic resistance, such as the acquisition/shuffling of resistance gene cassettes via the integron integrase, antibiotic resistance mutation via the error-prone polymerases, and potentially also antibiotic tolerance/persistence.

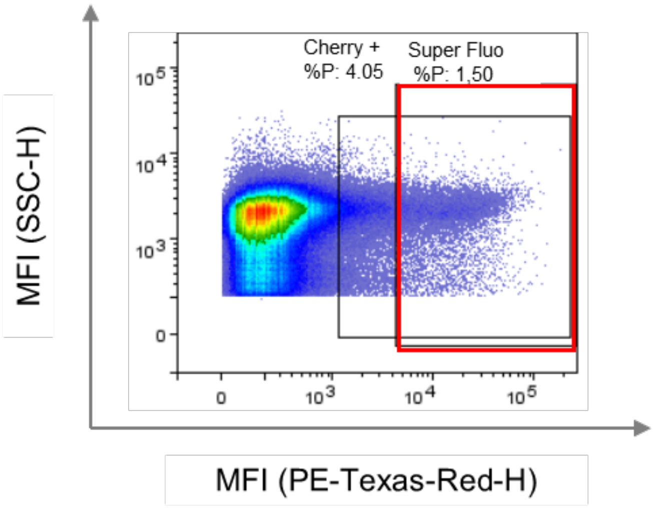


**Fig. S4: Quantification of cells expressing *sfiA* in biofilm grown from the sorted super-fluorescent population**

Sorted super-fluorescent population of strain MG F'/pPsfiA*cherry*-PintI1*intI1* was used to grow a biofilm in continuous culture in microfermentor for 24h at 37°C. The biofilm was analyzed by flow cytometry. Counting was performed after one hour of aerobic fluorescence recovery at room temperature. The dot-plot analyses is shown. The black square delimits the cherry-positive population (cherry +) based on the non -fluorescent strain MG F'/pSUΔtot and the red rectangle defines the *mcherry*-super-fluorescent population (super Fluo) based on derepressed *mcherry* expression in strain MG F'/ pPsfiA**cherry*-PintI1*intI1*. The data represent one representative experimental replicate.

### Table S1: Primers and Probes used in this study

| Construction | Primer name | Primer sequence | Reference |
|---|---|---|---|
| **MG1656λatt::*cherry*** | | | |
| | λ*att*.A1.500-5 | CGATGGCGATAATATTTCACC | [1] |
| | λ*att*.B1.500-3 | CCCTGATACTCACCAGGCATC | [1] |
| | λatt.A2.Amp-xfp-3 | GCGTTTTTTATTGGTGAGAATTACTAACTTGAGCGAAACG | [1] |
| | λatt.B2.Amp-xfp-5 | TGAGTAGGACAAATCCGCCGCTAAAAAGCAGGCTTCAACG | [1] |
| | λATT-ext5 | GGCGATAAATTGCCGCATCG | laboratory collection |
| | λATT-ext3 | TGCCACCATCAAGGGAAAGCCC | laboratory collection |
| | Kmfrt-XhoI-5 | AGCCTCGAGATGTGTGTAGGCTGGAGCTGCTTC | [1] |
| | Kmfrt-SacI-3 | CAGGAGCTCCTCATATGAATATCCTCCTTA | [1] |
| **pSU38Psfia*gfp+* and pSU38Psfia*gfp+*** | | | |
| | Psfia-gfp(+)INF-5 | CAGGGGCTGGATTGATTATGGATCCCAAAGGAGAAGAACTTTTC | This study |
| | Psfia-gfp(+)INF-3 | GCAAAAGCACCGCCGGACAAGCTTTTATTTGTAGAGCTCATCCATGC | This study |
| **pSU38Psfia*cherry* and pSU38Psfia*cherry*** | | | |
| | hindIII-mcherry | GATAAGCTTGTCGACTTACTTGT | This study |
| | bamHI-mcherry2 | CATGGATCCCAAGGGCGAGGAGGATAA | This study |
| **pSU38Psfia*cherry*-pintI*intI1*$_{\mathrm{w}}$ and pSU38Psfia*cherry-pintI*intI1*$_{\mathrm{w}}$** | | | |
| | ter-intI-fwd-2 | GAGCTCGTAAACTTGTCACCTCGAGGTCGACGG | This study |
| | inTI-pSU-rev | GGGTGTCAGTGAAGTACGCCTAGGTCTAGGGCGGC | This study |
| | ter-intI-rev-3 | CAAGTTTACGAGCTCGCTTGGAC | This study |
| | ter-ter-fwd-2 | GTAAACTTGGTCTGACGCTCAGTGGAACG | This study |
| | ter-ter-rev-2 | GTCAGACCAAGTTTACGAGCTCG | This study |
| | pSU-ter-fwd-2 | CCTGCCACATGAAGCGTCTGACGCTCAGTGGAACG | This study |
| **Sequencing** | | | |
| | IntI1 fwd | CGAACGCAGCGGTGGTAA | This study |
| | pZA for | GTCTTTCGACTGAGCCTTTC | laboratory collection |
| | MRV-D2 | TGCTGACGCACCGGTG | laboratory collection |
| | pSU38-verif-sq | GCCCACCCCCTGTCCCTC | laboratory collection |
| **qRT-PCR** | | | |
| | intI1-LC1 | GCCTTGATGTTACCCGAGAG | [2] |
| | intI1-LC5 | GATCGGTCGAATGCGTGT | [2] |
| | sfiA-L1 | TTACAGCAACTCGGTCAGCA | laboratory collection |
| | sfiA-R1 | CAGTGTGGCAAGGGGAGA | laboratory collection |
| | rpoB-R1 | GTTTGGTACGCGCAGAGAAG | This study |
| | rpoB-L2 | CCGGTATCGTTTACATTGGTG | This study |
| | cherry-L2 | GTGACCGTGACCCAGGACT | This study |
| | cherry-R1 | TGGTCTTCTTCTGCATTACGG | This study |
| | gyrB-L3 | GCTGCTGTTGACCTTCTTCTA | This study |
| | gyrB-R3 | TGTTCCTGCTTGCCTTTCT | This study |
| | kana R1 | GTAGCCGGATCAAGCGTATG | This study |
| | kana L1 | GAAGGGACTGGCTGCTATTG | This study |

| | | |
|---|---|---|
| sfiA-probe | [6FAM]CTGGGCTACCCTTAACGAAAGTAATGCAGA[TAM] | laboratory collection |
| rpoB-probe | [6FAM]CTGGTTGGTAAGGTAACGCCGAAAGGT [TAM] | laboratory collection |
| intI1-probe | [6FAM]ATTCCTGGCCGTGGTTCTGGGTTT [BHQ1] | [2] |
| gyrB-probe | [6FAM]TCGTCAGATGCCGGAAATCGTTGA [BHQ1] | This study |
| cherry-probe | [Cy5]CAGGACGGCGAGTTCATCTACAAGGT [BHQ2] | This study |
| kana-probe | [6FAM]TCCTGTCATCTCACCTTGCTCCTGC[BHQ1] | This study |


1. Lacotte, Y., Ploy, M.-C. & Raherison, S. Class 1 integrons are low-cost structures in Escherichia coli. *ISME J.* **11**, 1535–1544 (2017).
2. Barraud, O., Baclet, M. C., Denis, F. & Ploy, M. C. Quantitative multiplex real-time PCR for detecting class 1, 2 and 3 integrons. *J. Antimicrob. Chemother.* **65**, 1642–1645 (2010).
